# The matrisome contributes to the increased rigidity of the bovine ovarian cortex and provides a source of new bioengineering tools to investigate ovarian biology

**DOI:** 10.1101/2021.10.06.463107

**Authors:** Nathaniel F.C. Henning, Monica M. Laronda

## Abstract

The gonadotoxic effects of some cancers significantly increase the risk of developing infertility and cessation of ovary hormones (premature ovarian insufficiency, POI). Fertility preservation in the form of ovarian tissue cryopreservation (OTC) is offered to pediatric and adolescent cancer patients who cannot undergo oocyte retrieval and egg cryopreservation. The cryopreserved ovarian tissue can be transplanted back and has been found to restore fertility in 20 - 40% of transplants and restore hormone function for an average of 3 to 5 years. However, some individuals have primary or metastatic disease within their ovarian tissue and would not be able to transplant it back in its native form. Therefore, there is a need for additional methods for hormone and fertility restoration that would support a safe transplant with increased successful livebirths and long-term hormone restoration. To support these goal, we sought to understand the contribution of the ovarian microenvironment to its physical and biochemical properties to inform bioprosthetic ovary scaffolds that would support isolated follicles. Using atomic force microscopy (AFM), we determined that the bovine ovarian cortex was significantly more rigid than the medulla. To determine if this difference in rigidity was maintained in isolated matrisome proteins from bovine ovarian compartments, we cast and 3D printed hydrogels created from decellularized bovine ovarian cortex and medulla slices. The cast gels and 3D printed bioprosthetic ovary scaffolds from the cortex was still significantly more rigid than the medulla biomaterials. To expand our bioengineering toolbox that will aide in the investigation of how biochemical and physical cues may affect folliculogenesis, we sought to confirm the concentration of matrisome proteins in bovine ovarian compartments. The matrisome proteins, COL1, FN, EMILIN1 and AGRN were more abundant in the bovine ovarian cortex than the medulla. Whereas, VTN was more abundant in the medulla than the cortex and COL4 was present in similar amounts within both compartments. Finally, we removed proteins of interest, EMILIN1 and AGRN, from decellularized bovine ovarian cortex materials and confirmed that this specifically depleted these proteins without affecting the rigidity of cast or 3D printed hydrogels. Taken together our results indicate the existence of a rigidity gradient in the bovine ovary, that this rigidity gradient is maintained in resulting engineered materials strongly implicating a role for matrisome proteins in contributing to the physical properties of the bovine ovary. By establishing additional engineering tools we will continue to explore mechanisms behind matrisome-follicle interactions.

## 1. Introduction

In the U.S. more than 11,000 children between the ages of 0 to 14 are diagnosed with cancer each year.^1,2^ While treatments have dramatically increased the five-year survival rates of patients, with seven out of eight patients now surviving five or more years, the off-target effects of these treatments may cause long-term complications, such as premature ovarian inefficiency (POI).^3–7^ Longitudinal studies on POI patients have shown significant co-morbidities, which can include increased risk of cardiovascular disease and reduced bone health, resulting in a lifespan reduced by 2 years.^3–7^ There is only one fertility preservation option available to pediatric patients, ovarian tissue cryopreservation (OTC), in which the ovarian cortex is cryopreserved for future transplantation. While ovarian transplantation has resulted in more than 140 reported births only 20 - 40% of patient transplants restore fertility and the functional duration of the transplanted tissue, indicated by hormone restoration, is highly variable (range of 2 months - 12 years).^8–12^ Furthermore, studies have shown a risk of reintroducing disease in patients with ovarian metastasis via these transplants.^13^ Previous work has shown that a bioprosthetic ovary using isolated follicles in a 3D printed gelatin scaffold restores hormone production and fertility in ovariectomized mice which would allow us to isolate ovarian follicles from any metastatic cells.^14^ To improve upon this work and move toward translation for human use, a clear understanding of the ovarian microenvironment and how both physical and biochemical components of a transplantable scaffold can influence primordial follicle activation and subsequent depletion is needed. Indeed, human fertility restoration will require such improvements to scaffolds to support decades of normal function.

The ovary is compartmentalized into two regions, the cortex and the medulla. The outermost of these is the cortex containing the quiescent primordial follicles which make up the ovarian reserve while the innermost compartment, the medulla, contains most growing follicles. The ovarian extracellular matrix (ECM) can affect hormone availability and responsiveness by sequestering, trafficking or presenting factors including androgens and estrogens via sequestration of sex hormone binding globulin.^20,21^ Significant work has shown a role for the mechanical control of follicle activation and folliculogenesis using *in situ* primordial follicles or isolated growing follicles. These experiments revealed that primary follicles, found within the cortical region of the ovary, grow better when encapsulated in higher percentages of alginate while larger follicles, generally found in the medullary region, grow better in lower percentages.^22–29^ In fact, culturing a secondary follicle in a high percentage alginate results in transcriptional changes and increased androstenedione production.^30^ Additionally, exogenous physical pressure has been shown to maintain quiescence in primordial follicles that are released from their immediate environment, while disruption of the surrounding tissue induces increased primordial follicle activation via dysregulation of the HIPPO pathway.^30–34^ The composition and distribution of matrisome proteins were recently mapped across the porcine ovary. Forty-two matrisome proteins were significantly differentially expressed across the cortical and medullary compartments, revealing some proteins that may play critical roles in regulating primordial follicle activation and growth.^35^ Here we sought to further understand the physical properties of a mammalian ovary that contains distinct ovarian compartments and identify how the matrisome may contribute to the properties within these compartments. Furthermore, we created matrisome-based biomaterials and developed additional tools to explore the contribution of matrisome proteins, EMILIN1 and AGRN, found to be differentially distributed across ovarian compartments. This data and future functional assays will inform an improved bioprosthetic ovary.

## 2. Materials and Methods

### 2.1 Obtaining and Processing Bovine Ovaries

Bovine ovaries were purchased from Applied Reproductive Technology, LLC (ART, Wisconsin). Cows were post pubertal, but exact age ranges for the cattle at time of sacrifice and ovary retrieval was not available. After retrieval, ART washes the ovaries with 2% chlorohexidine gluconate diluted with distilled water followed by a series of washes with only distilled water. Ovaries were then shipped chilled overnight in phosphate buffered saline (PBS) with penicillin, streptomycin and gentamicin. Bovine ovaries without clear hemorrhagic cysts or other abnormal distortions were used for our experiments. For production of hydrogels, any large corpus lutea were removed prior to decellularization.

Bovine ovaries were placed in L-15 containing 1x antibiotic-antimycotic (Caisson, ABL02-100ML) solution prior to processing. Initial processing of tissue involved removing excess mesovarium and bisecting through the hilum. Tissue was further processed using a custom-made tissue slicer that produces slices that are 1 mm thick (Northwestern Simulation Lab). Tissue was sliced on the cut side after bisecting to allow atomic force microscopy (AFM) sampling from both cortex (first 0.5 mm) and medullary regions (>0.5 mm). Tissue slices were then placed in DMEM containing 1x antibiotic-antimycotic overnight at 4°C.

### 2.2 Atomic Force Microscopy for Mapping Ovarian Rigidity

AFM was carried out using a Bruker Hysitron BioSoft Indenter with a 100 μM radius probe. Bruker Bioscan (v1.0.0.1) software was used for data acquisition and Origin 2018 software was used for data analysis and force curve fitting to a Hertzian model as recommended by core facility. Prior to AFM testing, 1 mm thick slices of ovarian tissue were rinsed with PBS (Thermo, 10010023), then adhered to 35 mm petri dishes using PELCO Pro CA44 Instant Tissue Adhesive (Pelco, 10033). Samples were then submerged in PBS to prevent drying. The AFM probe was calibrated in PBS to remove background noise from the liquid interface. This probe was manually brought to the surface of the bovine ovary slice. Once the surface was reached, the probe was manually retracted from the surface prior to data acquisition. Data acquisition was performed using a three-step protocol (approach, hold, retraction) with the following settings: approach/retraction rate of 5.00 μM/s with a target value of 150 μm, and a 5 seconds hold.

### 2.3 Decellularization

Decellularization was carried out using 0.1 % sodium dodecyl sulfate (SDS, Sigma, 75746) in PBS (Thermo, 10010023) as used previously.^35^ Slices of tissue and isolated medulla were placed on a nutator at 4 °C and SDS solution was changed every 24 hours for 48 - 72 hours for slices or 2 - 3 weeks for larger medulla pieces prior to use in ink manufacturing.

### 2.4 Ovarian Hydrogel Manufacturing

Decellularized ovarian tissues were lyophilized using a Freezone 6 Plus System. After lyophilization, tissue was milled until it fit through a 60-mesh screen. Residual SDS and lipids, which would interfere with gelation, were subsequently removed using three 100% ethanol washes and the washed milled tissue was subsequently lyophilized again using the Freezone 6 Plus System. Pepsin digestion is then performed on the tissue, 20 mL of a 1 mg/ml pepsin (Sigma P7012) digest solution in 0.1 M HCl. The pH of the solution was verified to be ~1-2 (the ideal range for pepsin activity) and then 25 mg/mL of tissue was added and placed on a magnetic stir plate for 72 - 96 hours at room temperature. After digestion, the solution was neutralized using NaOH. The gel was then placed at 37°C for 1 hour prior to use to promote gelation in either 3D printing or as a cast gel.

### 2.5 3D Printing

The 3D printed scaffold design was used previously by Laronda, et al.^11^ The design is a 15 mm x 15 mm square that includes a solid bottom, with each subsequent layer printed at a 60° advancing angle and struts were spaced 1 mm apart. Five layers were printed using this design for use with AFM-based analysis. FRESH printing was performed as described previously.^36^ Cortex and medulla-derived dECM inks were mixed 1:1 with high concentration COL1 Lifeink 200 (Advanced Biomatrix, 5278) to assist with printability (COL1 control data included in Supplementary Figure 3). Hydrogel was loaded into low temperature cartridges for use with an Envisiontec 3D-Bioblotter manufacturing model. After mixing and loading into cartridges they were incubated in the print head to obtain a uniform temperature of 8 °C, then printed using 32-gauge needles (Nordson, 7018462).

### 2.6 Atomic Force Microscopy for Engineered Materials

AFM was carried out on engineered materials using a Piuma Nanoindenter with accompanying software (V3.2.0). Prior to AFM runs, printed scaffolds were rinsed with PBS then adhered to 35 mm petri dishes using PELCO Pro CA44 Instant Tissue Adhesive (Pelco, 10033). Subsequently, samples were submerged in PBS to prevent scaffolds from drying out. For cast gels, a small volume of gel was added to a transwell insert (Millipore, PICM01250). Cast gels were then spun in a centrifuge for 5 minutes at 2,000 xg to remove bubbles. After centrifugation a small volume of PBS was added to the top of the gels to assist with AFM measurements. Analyses for all materials was carried out using a probe manufactured by Optics 11 with the following properties: a rigidity of 0.033 N/m rigidity, and a tip radius of 29 μM. The AFM probe was calibrated in PBS to remove background noise from the liquid interface and was calibrated against a plastic dish as per Piuma calibration requirements. Piuma software was used to find the surface of materials and then to acquire data. Data acquisition was carried out using a three-step protocol (approach, hold, retraction) with the following settings: An approach/retraction rate of 5.00 μM/s with a target value of 150 μm, and a hold time of 5 seconds.

### 2.7 Quantitative iPCR

Protein was extracted from bovine ovary slices that were decellularized (as described above). Tissue was placed in a protein extraction buffer made of 1% SDS, 50 mM ammonium bicarbonate, 50 mM NaCl, and 10 μL/mL Halt Protease Inhibitors (Cell Signaling Technology, 5872S). Tissue was placed in reinforced 2 mL tubes (Omni International, 19–648) with 2.8 mm polycarbonate beads (Omni International, 19–646). Tissue was homogenized with an Omni BeadRuptor12 at 4 °C (Omni International, 19-050 A) using the following settings: 6 cycles at speed 6.0, 45 second homogenization and a 75 second delay between cycles. Homogenates were subsequently sonicated on ice 3 times at 75% amplification for 1 minute. Samples were then centrifuged at 10,000 rpm for 15 minutes. A 1 mL aliquot was taken from the samples and samples were treated with an SDS-Out kit (Thermo Scientific, 20308). Protein concentration was measured using a BCA assay (Fisher Scientific, PI23227). Samples were normalized to a concentration of 1,000 ng/μL using protein lysate buffer from the Taqman Open Kit (Thermo Fisher, 4453745).

Antibodies were biotinylated using EZ-Link Sulfo-NHS-LC-Biotin, No-Weight format kit (ThermoFisher, 21327). Antibody information is listed in Supplementary Table 1. Excess biotin was removed using two cycles of filtration with Zeba Micro Spin Desalting columns, 40k (Fisher Scientific, PI87765). Probes were then tested for suitability using the method described in the Taqman Protein Assays Probe Development Protocol (Thermo Fisher, 4448549). To compare protein quantity between the medulla and cortex a standard method described in the Taqman Protein Assays Sample Prep and Assay (Thermo Fisher, 4453745) was used. A four-point dilution series was created with 2000 ng of total protein then serially diluted 1:10. A qPCR machine (Applied Biosystems, QuantStudio 3) was used for running the assay. To quantify the amount of protein of interest present a standard curve was created with proteins of known concentrations in a dilution series (see Supplementary Table 1). Because some samples had undetectable levels of protein, dose curve data are presented as change in CT over no template controls (see Supplementary Figure 1). Two-way ANOVA and multiple t-test analyses were used to determine significance, GraphPad PRISM version 9 was used to interpolate concentrations based on the standard curves.

### 2.8 Targeted Depletion of Matrisome Proteins

Targeted depletion of gels was carried out using antibodies for the protein of interest and a seven-step magnetic assisted depletion process. Antibodies used for targeted depletion were prepared using goat anti-rabbit magnetic beads (NEB, S1432S) following the manufacturer’s protocol. Tagged antibody was added to decellularized extracellular matrix (dECM) homogenate following manufacturers recommended concentration in a 1.5 mL Eppendorf tube. After incubation with the antibody a strong magnet was applied and the dECM was transferred to the next tube for further depletion. This process was repeated 7 times to maximize reduction.

## 2. Results and Discussion

### 2.1 AFM measurements revealed a rigidity gradient in the bovine ovary that can be recapitulated in engineered materials

#### There is a rigidity gradient across in bovine ovarian compartments

The rigidity of the bovine ovary was measured using AFM nanoindentation. Young’s moduli of bovine ovary slices were measured at points spaced in 1.0 mm increments from the ovarian surface epithelium inward, where 0.5 mm from the ovarian surface epithelium edge at the mesosalpinx plane was considered the cortical region and 1.5 – 3.5 mm points inward along the central pole were considered the medullary region (**Figure 1A).** A tissue thickness of 1 mm was chosen to eliminate possible contributions from the 35 mm dish to rigidity measurements being taken by the AFM. Further, a probe size of 100 μm was used to better detect the overall rigidity of a region, and to avoid the over-representation of small ovarian structures in rigidity measurements. Two technical replicates were taken per ovary slice, with 4 - 5 biological replicates taken for each distance. There was a statistically significant difference between the rigidity of the cortex and the medulla (**Figure 1A**). The rigidity of the ovary was significantly reduced from the cortex (8.87 kPa) to 3.5 mm (1.05 kPa). The rigidity values ranged in the ovary from 0.71 to 11.59 kPa. Overall, the cortex was 8.5 times more rigid than the deep medulla (3.5 mm) and the deep medulla was approximately two-fold (average 2.05) less rigid per 1 mm step as measured by AFM.

**Figure 1.**
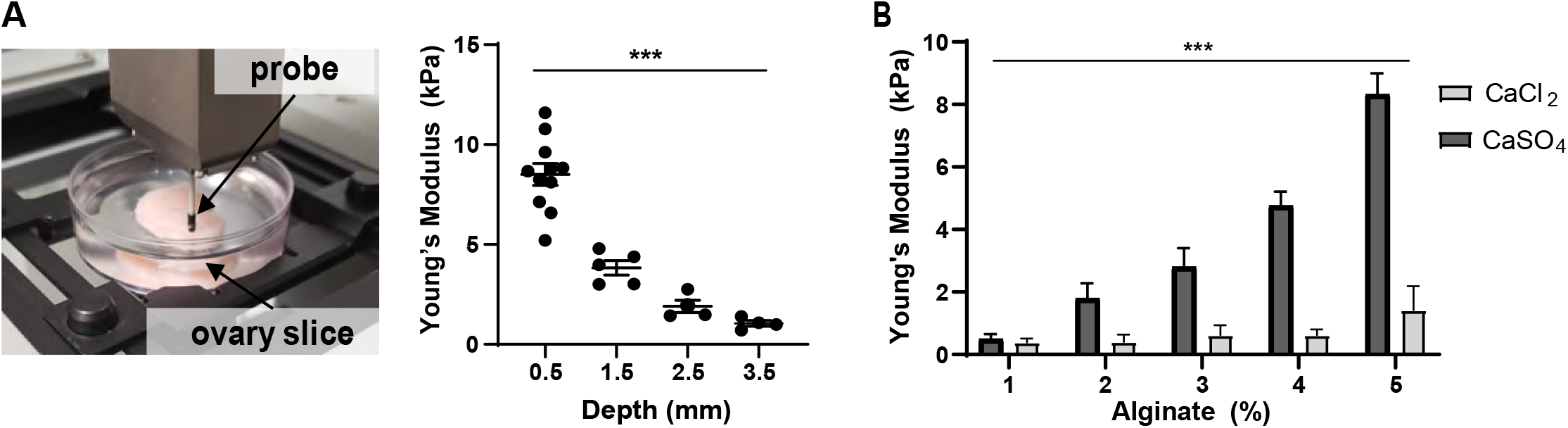
Rigidity of Bovine Ovary and Materials Used in Ovarian Culture. (A) AFM was used to measure the rigidity of bovine ovary tissue (left) at 0.5 mm to 3.5 mm from ovarian surface epithelium inward (right). p-value = <0.0001 (B) AFM was also used to measure the rigidity of cast alginate gels cross-linked with either CaCl_2_ or CaSO_4_.

#### CaCl_2_ crosslinked alginate does not recapitulate bovine cortex rigidity

Several seminal studies on follicular growth, maturation, and transcriptional changes were performed with different percentages of alginate with most papers focusing on alginate gels between 0.5%-3% with 0.5-1.5% alginate being used as permissive environments for culturing growing follicles and 2-3% being used to represent rigid environments for culturing primary follicles.^22–29,80^ However, previous investigations of the rigidity of this material at different percentages were performed using different techniques, specifically rheometry.^80^ Therefore, we sought to define the rigidity of alginate using AFM in order to compare it to the native environment described above. We first examined the rigidities of alginate gels using varying percentages of alginate cross-linked with either CaCl_2_, the most common cross-linking substrate used in follicle culture (**Figure 1B**).^22–29^ CaSO_4_ was chosen as a second cross-linker as previous research has shown that alginate gels cross-linked with CaSO_4_ achieved higher rigidities than those crosslinked with CaCl_2_, yet was still able to support murine follicle survival in culture.^36^ The rigidity of alginate crosslinked with CaCl_2_ ranged from 0.39 ± 0.13 kPa to 1.42 ± 0.77 kPa, which was significantly less than alginate crosslinked with CaSO_4_ which ranged from 0.50 ± 0.15 kPa to 8.33 ± 0.67kPa. Measurements for CaCl_2_ crosslinked gels failed to reach higher rigidities found in bovine ovaries even at the maximum alginate percentage tested (5%, 1.42 kPa). However, CaSO_4_ crosslinked gels showed similar rigidities to bovine ovaries at 5% (8.33 kPa) and 2% (1.80 kPa) being equivalent to rigidities at the cortex (8.87 kPa) and the deep medulla (1.05 kPa) respectively. In comparison 5% CaCl_2_ crosslinked gels, the maximum percentage tested, only showed similar rigidity to the deep medulla.

#### Engineered materials using decellularized bovine ovaries recapitulate compartmental rigidity differences

In order to examine compartment-specific contributions to rigidity from the matrisome, we performed AFM analysis on both cast hydrogels and 3D printed scaffolds created from decellularized bovine ovary material. Slices from the cortex and medulla were decellularized and lyophilized. The enriched matrisome material was then milled, washed, re-lyophilized and digested with pepsin to create compartment specific hydrogels (**Figure 2A)**. First, the rigidity of cast gels were analyzed using AFM (**Figure 2B)**. Cortex gels had a rigidity of 3.11 kPa ± 0.41, which was significantly higher than gels derived from the medulla material, which were 0.02 kPa ± 0.001. The gel made from the cortex maintains the higher rigidity over the medulla and is 155.5 times more rigid. The cortical matrisome gel is 2.85 times less rigid than the native cortical region (0.5 mm depth); whereas the medulla matrisome gel is 112 times less rigid than the native medullary compartment (1.5 – 3.5 mm). Then, to determine if these hydrogels maintained their rigidity differences within printed scaffolds, we 3D printed scaffolds in a similar architectural design as previously demonstrated to support murine ovarian follicle growth and maturation.^11^ To increase printability, the hydrogels derived from cortex and medulla were mixed 1:1 with collagen 1 (COL1) ink then printed using FRESH methods prior to examining rigidity through AFM (**Figure 2C**).^36^ FRESH printed scaffolds containing cortical matrisome were significantly more rigid than those printed with medullary matrisome materials with average rigidities of 4.99 kPa ± 1.52 and 0.06 kPa ± 0.002, respectively. This pattern of cortical materials measuring significantly higher rigidities than medulla materials was maintained even in printed scaffolds. The cortex scaffolds were 83 times more rigid than those derived from the medulla. AFM analysis of scaffolds using only COL1 ink had an average rigidity of 0.17 kPa ± 0.05, which was significantly less than the cortex-derived scaffolds and trending toward statistically significantly higher than medulla-derived scaffolds (p = 0.078, **Supplementary Figure 2**).

**Figure 2.**
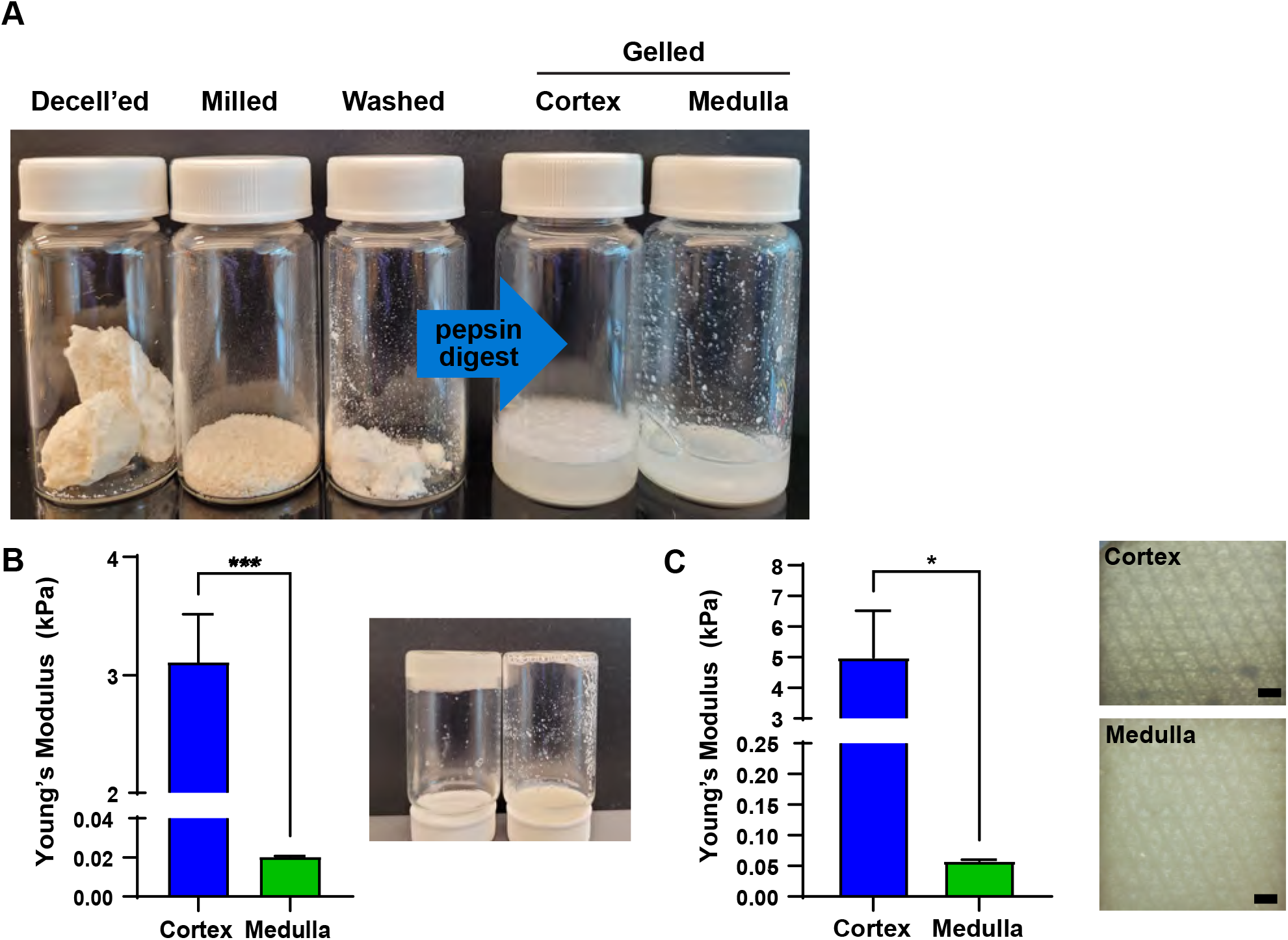
Mechanical analysis of Engineered Materials Using AFM. (A) Ovary slices from cortex or medulla (shown) were decellularized (decell’ed) and lyophilized; then milled, washed and re-lyophilized before being digested using pepsin to make gels. (B) The rigidity of cast gels from cortex or medulla material was measured with AFM. The cortex made stronger gel than the medulla (right). p = 0.0003 (C) The rigidity of scaffolds printed from cortex and medulla hydrogels mixed with collagen in 1:1 volume was measured using AFM. p = 0.017, scale = 200 μm

### 2.2 Quantitative iPCR revealed differential distribution of matrisome proteins across ovarian compartments in bovine ovaries

To examine the distribution of matrisome proteins across ovarian compartments we performed iPCR on slices of bovine ovaries that were decellularized to enrich for matrisome proteins. Consistent with the above methods, we considered the first 0.5 mm slice cortex, while the rest of the ovary was considered medulla. Candidate proteins for this analysis were chosen based on previous experiments in porcine ovaries.^15^ We chose the following candidates: COL1 and collagen IV (COL4), agrin (AGRN), elastin microfibril interfacer 1 (EMILIN1), fibronectin (FN1), and vitronectin (VTN). To quantify these proteins a standard curve was created with purified protein (see **Supplementary Table 1**). There were significant differences in the distribution of candidate proteins across ovarian compartments that matched the previously published mass spectrometry and iPCR analyses from the porcine ovarian matrisome map (**Figure 3**). COL1, AGRN, EMILIN1, and FN1 are significantly more abundant in the cortex matrisome than the medulla. Whereas, VTN is significantly more abundant in the medulla and the amount of COL4 is not different among the two compartments. This was the first time iPCR has been used to quantify matrisome proteins in tissues. Two proteins of interest, EMILIN1, a matrisome glycoprotein, and AGRN, a matrisome proteoglycan, have been shown to act upstream of proliferation and mechanotransduction pathways in other tissues. ^37–48^

**Figure 3.**
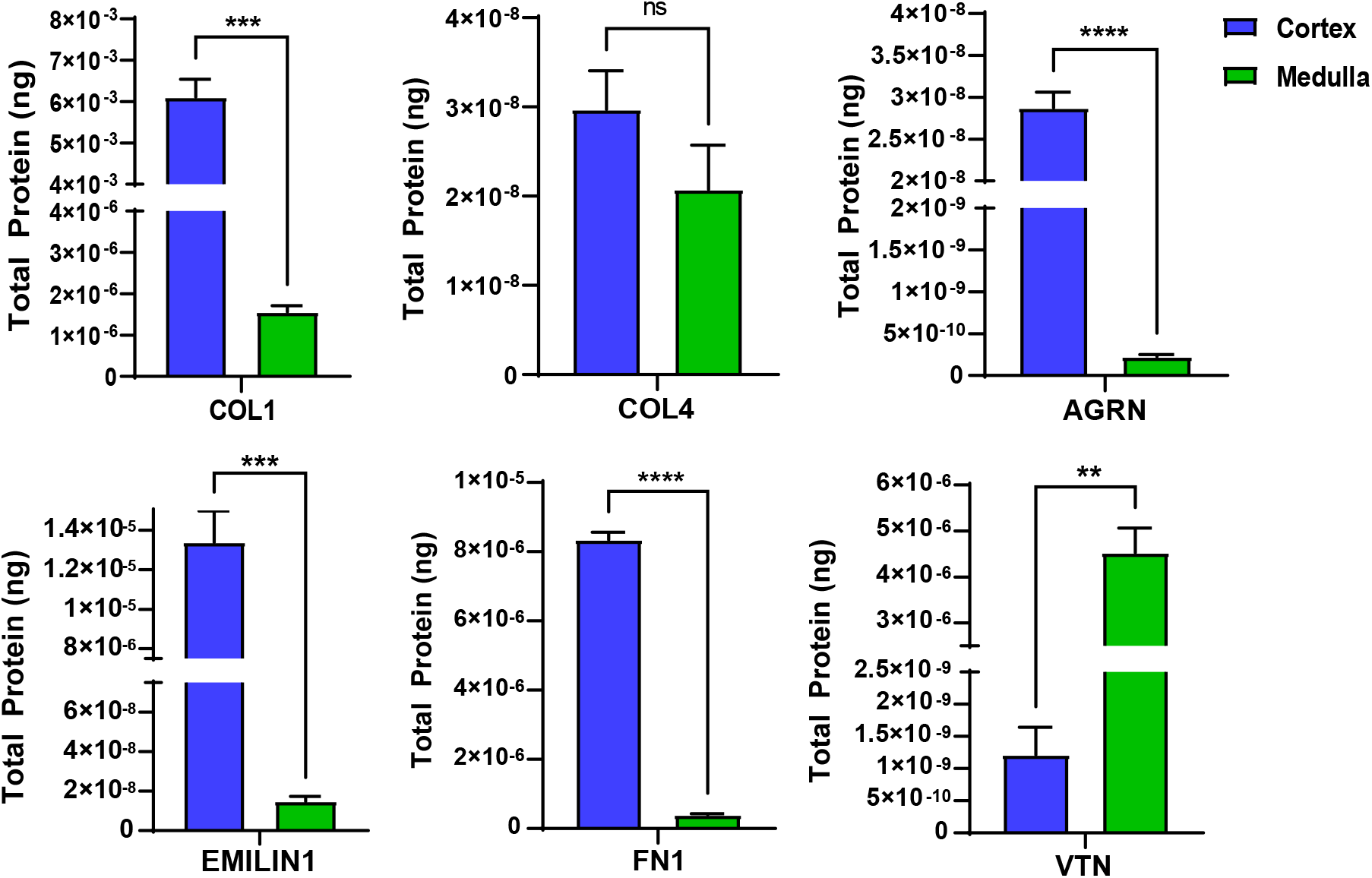
Quantitative iPCR analysis of matrisome proteins. Total COL1, COL4, AGRN, EMILIN1, FN1 and VTN protein within decellularized bovine ovary cortex and medulla. Bars represent mean with standard error. ns, not significant, ** = 0.001, *** < 0.001, **** < 0.0001.

### 2.3 Targeted depletion of matrisome proteins from engineered materials

#### Magnet assisted protein filtration specifically depletes proteins of interest in tissue matrisome samples

In order to explore the effects or matrisome proteins on follicle activation and folliculogenesis we developed a novel toolkit to study these mechanistic relationships by selectively depleting proteins of interest using magnet assisted protein filtration (MAPF) from ovarian tissue derived matrisome samples. For the purposes of this study we selectively depleted and removed EMILIN1 and AGRN. Cortex matrisome materials were used because of the significant abundance of these proteins in this compartment and iPCR was used to measure the remaining proteins of interest against the percentage of an abundant off-target protein, COL1. We determined that a seven-step filtration process significantly reduced the amount of both EMILIN1 and AGRN from cortical matrisome materials to satisfy future knockdown experiments (**Figure 4A, 4B)**. Indeed, the seven-step filtration method reduced EMILIN1 by 97.714% ± 7.89 and AGRN by 96.80% ± 19.11 while not affecting the amount COL1 (102% ± 22.76 for EMILIN1 targeted filtration, and 95.36% ± 4.89 for AGRN) that was present in the same samples.

**Figure 4.**
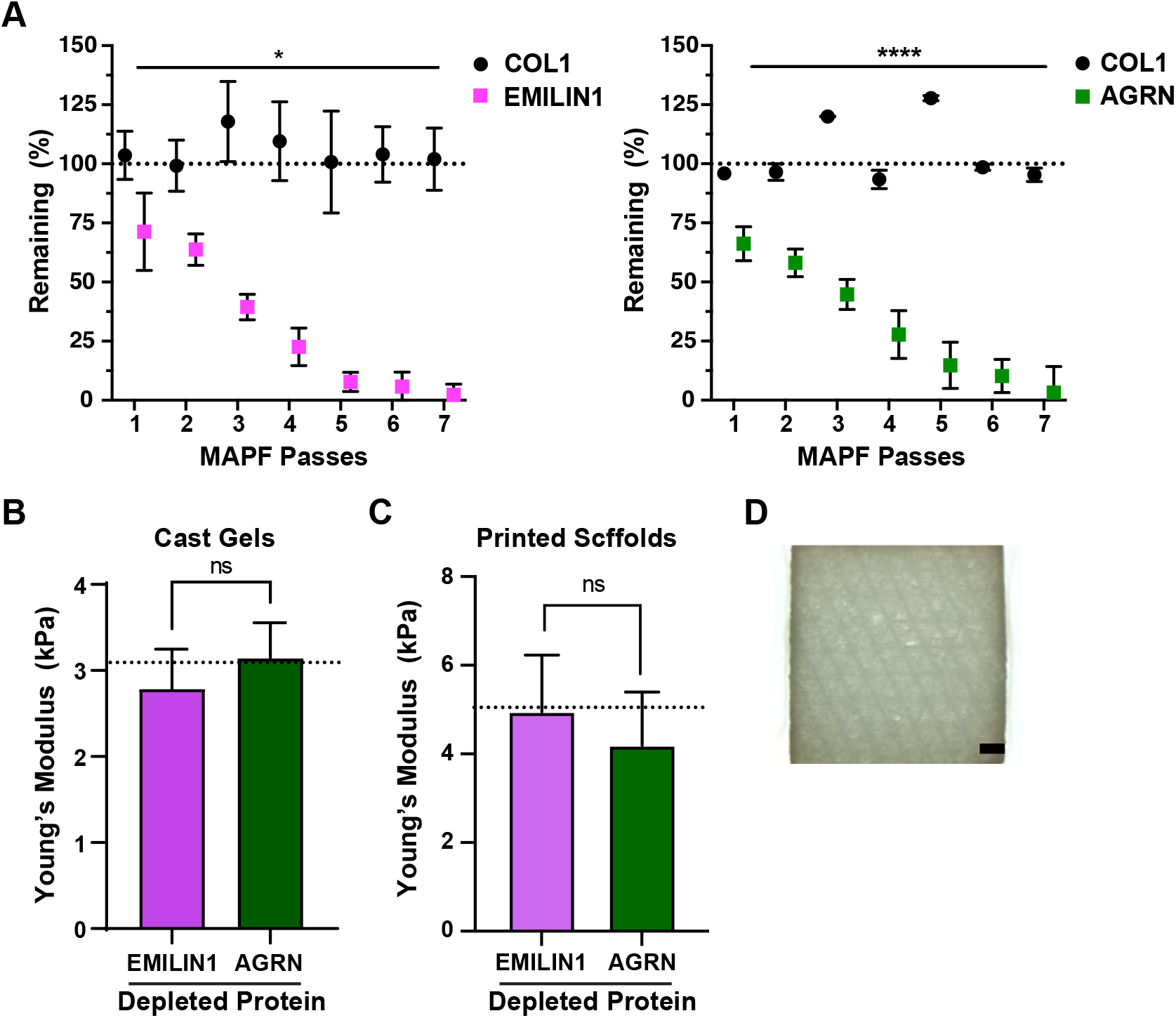
Targeted depletion of proteins of interest and mechanical analysis of resulting hydrogels. (A) iPCR analysis of relative content of COL1 and EMILIN1 or AGRN following targetted depletion of EMILIN1 or AGRN by passing cortical matrisome samples through magnetic activated protein filtration (MAPF). Dotted line = 100 % (B) Rigidity of cast or (C) printed gels with EMILIN1 or AGRN depleted from cortical matrisome materials by seven MAPF passes. Dotted lines = rigidity of control cortex gels (B) or cortex printed scaffolds (C). (D) Light microscopy image of scaffold printed with cortex matrisome (top) and depleted cortex matrisome (bottom). Bars, squares or dots represent mean with standard error. ns, not significant, * = 0.01, **** < 0.0001, scale = 200 μm

#### Physical properties of selectively depleted materials are unaffected

In order to assess the physical properties of matrisome materials that underwent the MAPF process we tested the rigidity of cast gels and printed scaffolds. There were no significant differences in the rigidity as measured by AFM of depleted gels (**Figure 4C)** or scaffolds (**Figure 4D**) in comparison to the base hydrogels from cortical matrisome materials. This indicates that the MAPF process and the individual removal of these matrisome proteins does not affect the rigidity of engineered materials, which is a positive indicator for its use as an *in vitro* platform to explore the influence of matrisome proteins on folliculogenesis.

## 3. Discussion

In this study we examined the rigidity of bovine ovaries, assessed ovarian compartment specific matrisome hydrogels, and expanded our toolkit to examine the mechanistic connection between the matrisome and folliculogenesis. We found that the bovine ovary contains a rigidity gradient with a cortex that is significantly more rigid than that of the medulla, that this difference in rigidity is maintained in engineered materials (cast gels and printed scaffolds) derived from these compartments, and that we can use MAPF-based methods to deplete proteins of interest. We used the same AFM technique to measure the rigidity of encapsulation materials commonly used for *in vitro* folliculogenesis experiments to compare these measurements with native and decellularized bovine ovary materials. Additionally, we confirmed that the bovine ovary contains the same profile for COL1, COL4, AGRN, EMILIN1, FN1 and VTN as previously demonstrated in porcine ovaries. For the first time, we quantified the amount of protein in each sample using iPCR. This technique may be useful in building an engineered environment that matches the protein abundance of native tissue. Finally, the new hydrogel materials, cortical and medullary hydrogels and hydrogels with depleted proteins of interest expand our toolkit to facilitate future investigations, including into molecular mechanisms that begin with external matrisome proteins.

Using AFM we showed that the bovine ovary has a rigidity that ranges from 0.71 to 11.59 kPa. To place this within context of other organ systems, the rigidity of the ovary contains rigidities seen within many other soft tissue organs (muscle, lung, liver, and kidney, and brain).^49–54^ Our findings demonstrated that a rigidity gradient exists between ovarian compartments, not just in native tissue but also within hydrogels formed from matrisome proteins isolated from different compartments. These findings support the hypothesis that primordial follicles, located within the cortex, reside within a more rigid environment than activated or growing follicles within the medulla.^81,82^ The significant rigidity differences between compartments also support the growing body of research that implicates multiple mechanotransduction pathways as playing a role in natural and induced folliculogenesis.^32,60–64^ Kawamura, et al in particular showed increases in follicle activation after cutting the ovary with a subsequent disruption of Hippo signaling stimulating follicle growth.^32^

The data here, however, conflicts with two recent studies, one examining ultrasound shear wave velocities in bovine ovaries and another which examined the micromechanical properties of the murine ovary.^65,66^ Both studies indicated that the medullary region was more rigid than the cortex with the study by Hopkins, et al concluding that the major driver of rigidity changes in the murine ovary is most closely associated with mature follicles. However, these differences in bovine versus murine ovary rigidity measurements using the same nanoindentation methods could indicate that significant species differences exist in the physical properties and organization of the ovary. These differences may be a result of differences in structure between bovine (similar to the human ovary) and murine ovaries that can be specifically identified during organogenesis where less regionalization develops in the murine ovary compared to the human ovary.^68^ The research by Gargus, et al examined whole bovine ovaries through shear wave velocity.^66^ We chose to use nanoindentation methods over other methods that measure viscoelastic properties because nanoindentation can work on the micron scale. However, alterations in shear wave velocity could indicate other changes across compartments associated with differences in ECM architecture and composition, density, size, and shape of follicles and other structures.^69,70^ One limitation of the study here is that we processed the ovary into 1 mm slices and disrupted existing structures within the ovary that would have surface tension *in vivo*.

In order to expand our toolbox to include biomaterials that could be used to better understand the role of the microenvironment on folliculogenesis, we examined gels and inks made from decellularized cortical and medullary slices from bovine ovaries. Using a consistent method to define rigidity of these materials and to examine the contribution of matrisome proteins to the rigidity of ovary was key. Our data complemented recent work by Amargant, et al, that demonstrated, using AFM, that the increase in rigidity of aged murine ovaries could be contributed to the increased COL1 content indicative of fibrosis.^67^ The data presented here may offer some explanation as to why primordial follicles from multiple species (primate, mouse, and human) survive and grow better in a higher percent alginate than in the more permissive alginate that is preferred by growing follicles. ^22–29,55^ To compare the rigidity of these commonly used materials with the native tissue and newly created biomaterials, we performed AFM on alginate gels. Notably, we found that the highest rigidity gels used in previous experiments (3% alginate crosslinked with CaCl_2_) wouldn’t recapitulate the rigidities seen in the cortex of native bovine ovaries.^80^ However, culturing a growing secondary follicle in a high percentage alginate results in transcriptional changes and increased the ratio of androstenedione to estradiol produced which can may be representative of a diseased state.^56–59^ While our goal is to create a scaffold or hydrogel microenvironment that mimics the physical and biochemical cues of the native ovary, we cannot overlook the contribution of the interstitial and neighboring follicles to the rigidity in the native ovary. Indeed, the matrisome gels were less rigid than native tissues (1.78 times less for 3D printed scaffolds, 2.85 for cast gels). These differences between native bovine ovary rigidity and the rigidity of biomaterials may also be attributed to the organization of matrisome proteins and fibers. These are properties that may be lost during the decellularization and homogenization methods required to produce matrisome hydrogels. Support of this comes from current efforts for scaffold design examining fiber morphology, ECM network architecture, and topography in addition to viscoelastic properties such as those defined by AFM analysis.^72–75^

Finally, we developed a novel toolkit for examining the contribution of matrisome proteins to follicle activation and folliculogenesis by using MAPF to deplete proteins of interest from engineered materials without disrupting the physical properties of resulting scaffolds or cast gels. The reduction seen using MAPF is comparable to other methods for selective depletion of proteins that are currently in use. ^77–79^ These EMILIN1 and AGRN depleted hydrogels are additional tools that will assist in isolating the role of biochemical cues from physical cues. One possible conclusion that could be drawn from these results is that neither AGRN nor EMILIN1 are directly associated with the physical properties of the ovary. Intriguingly, previous work done in EMILIN1 -/- mice has indicated that these mice show increased rigidity of the aorta in both nano and macro scale experiments.^76^ The lack of change in the engineered materials utilized here may indicate that EMILIN1 is primarily a driver of a cellular response leading to increased fibrosis, secretion of ECM components, etc. instead of organizing the ECM. Further work will need to be done to explore if the reduction is physiologically relevant to pathways of interest or if the remaining protein is capable of reaching a critical threshold needed for signaling in biological systems.

## 4. Conclusion

We have for the first time examined, quantified, and mapped the rigidity of the bovine ovaries across the anatomical compartments using AFM. Our data reveals that the bovine ovary has a range of rigidity of 1.4 – 8.86 kPa. This data supports the previous observation that the cortex is more rigid than the medulla with a statistically significant decline in rigidity from the outer edge of the ovary toward the center. Then we found that compartmental rigidity of hydrogels and 3D printed scaffolds made from the cortical or medullary compartments of decellularized ovaries retain the physical properties of their originating compartment with cortex derived materials being more rigid than medulla derived materials. Additionally, for the first time, we quantified the content of matrisome proteins in a biological material using iPCR. This revealed that COL1, COL4, AGRN, EMILIN1, FN1 and VTN matched the pattern previously identified in an unbiased matrisome analysis across compartments of the porcine ovary^35^. Finally, we developed a toolkit for examining the contribution of matrisome proteins to follicle activation and folliculogenesis by selectively depleting the candidate proteins EMILIN1 and AGRN using MAPF. We demonstrated that selective depletion of EMILIN1 and AGRN did not alter the rigidity of gels or printed scaffolds in comparison to undepleted hydrogels and inks. This may not be true for all proteins, and it is likely that some selective depletion of fibrous proteins would affect the rigidity of the resultant biomaterial products. We predict that this foundational work will support future work that will inform the development of bioinks for 3D printed scaffolds by allowing for the incorporation of important physical and biochemical cues that yield the desired follicular behavior. This research enables future implementation of an effective scaffold for a bioprosthetic ovary transplant.

## ACKNOWLEDGMENTS

This work is supported by the Warren and Eloise Batts Endowment (MML), Burroughs Wellcome Fund Career Award at the Scientific Interface (MML), and NIH/NICHD R01HD104683. AFM analysis was performed in the Analytical bioNanoTechnology Core Facility of the Simpson Querrey Institute at Northwestern University. ANTEC is currently supported by the Soft and Hybrid Nanotechnology Experimental (SHyNE) Resource (NSF ECCS-2025633). The Innovations lab in the Northwestern Simulation program was used to produce the custom 1 mm tissue slicer. We thank Dr. Adam Feinberg and his lab at Carnegie-Mellon University, Pittsburgh, PA, especially Maria Stang and Dr. Feinberg, for generously providing the FRESH materials and providing invaluable insight into printing our organ-derived matrisome protein inks.

## DATA AVAILABILITY

The raw data required to reproduce these findings are available upon request to the corresponding author.

---

**Supplementary Table 1:**
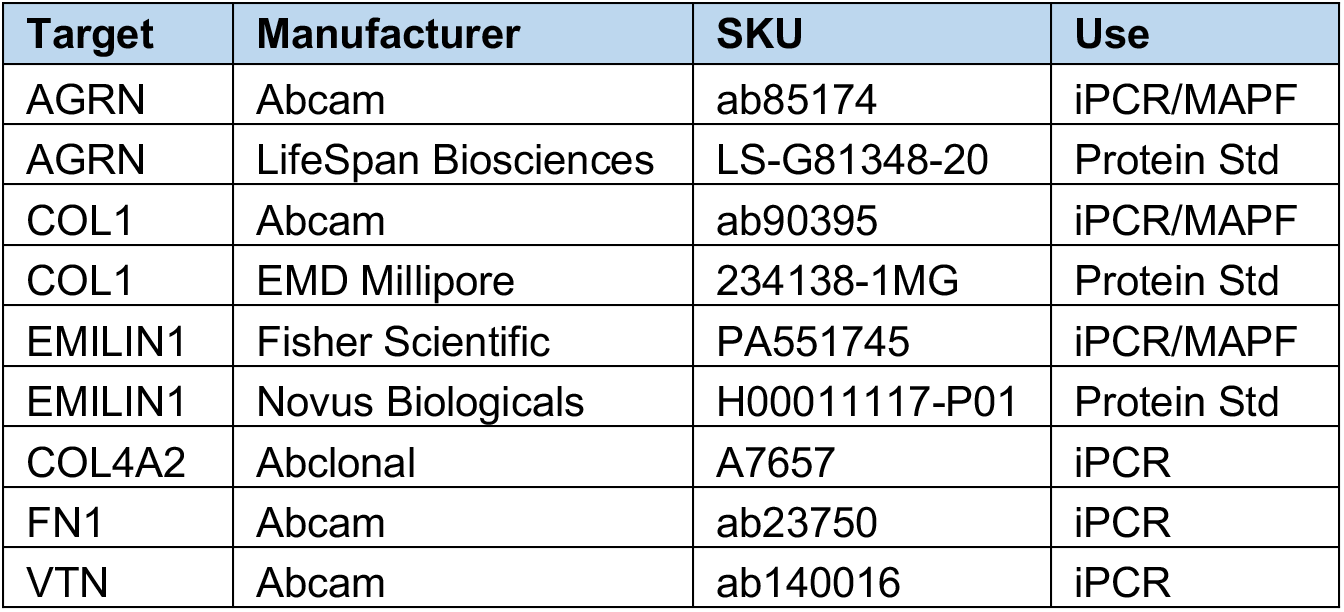
Antibodies and Proteins used for iPCR and MAPF

**Supplementary Figure 1.**
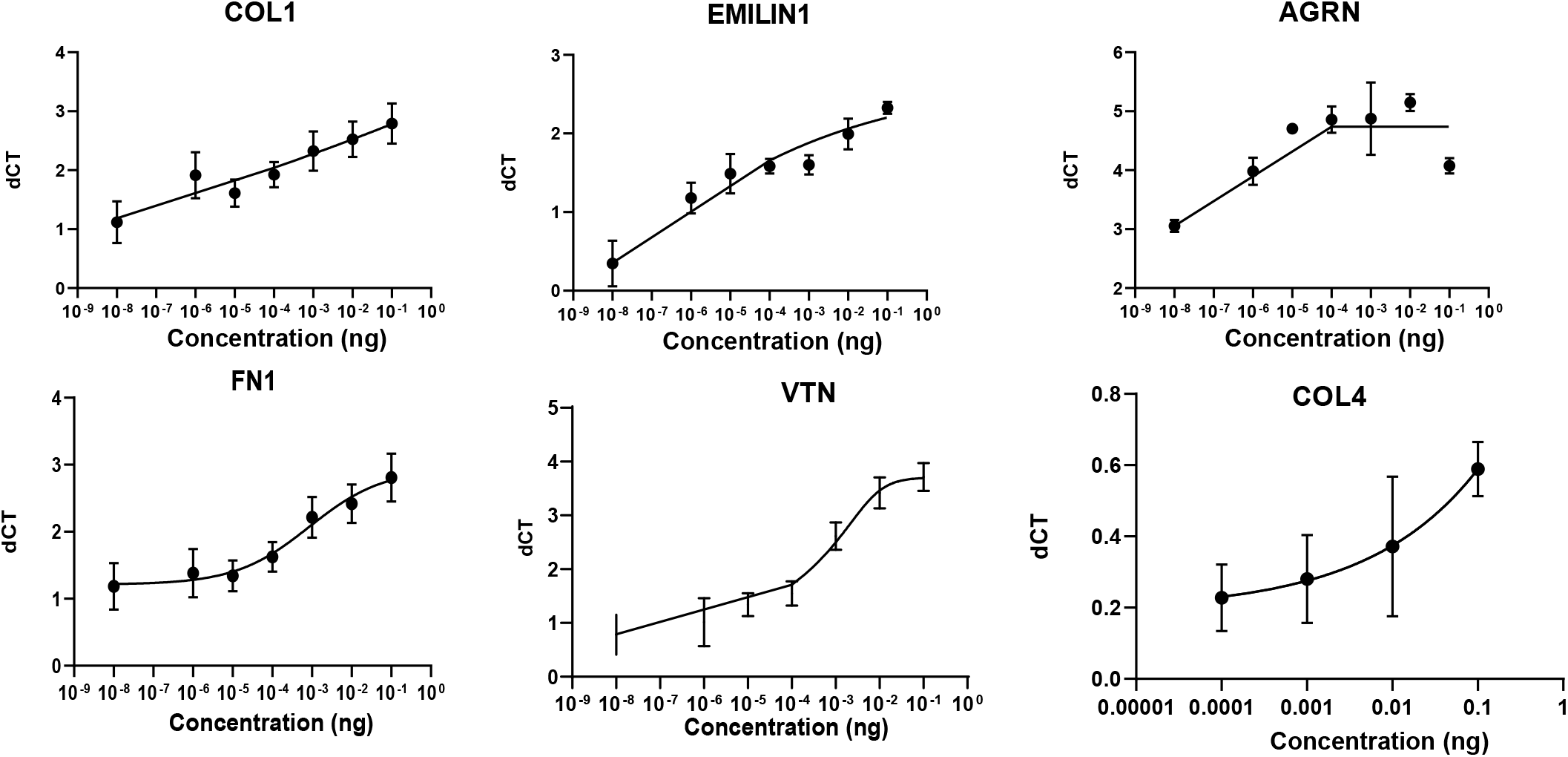
Standard curves used for interpolation of concentrations of protein. Bars represented as mean, SEM; N = 3 biological replicates, 2 technical replicates.

**Supplementary Figure 2.**
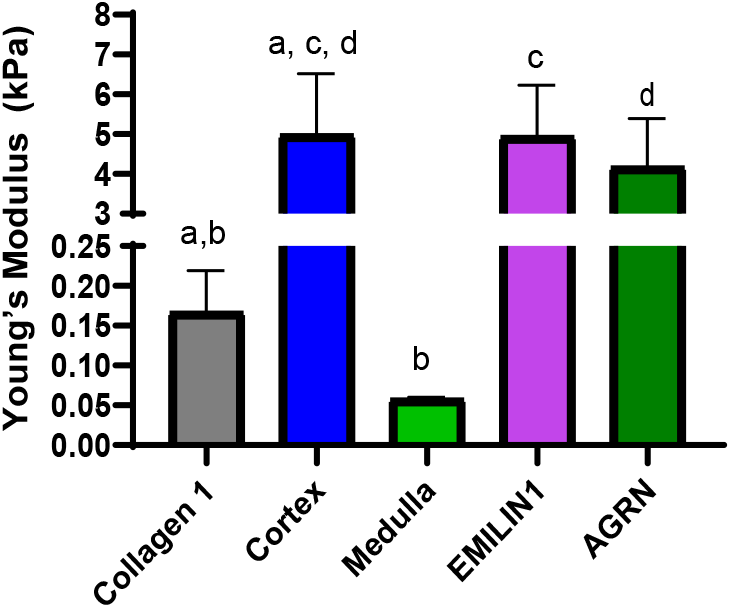
Combined AFM data for FRESH printed scaffolds including collagen 1 only scaffold control. Bars equal to mean and standard error, P-values for relationships: a=0.0196, b=0.0782, c=0.9825, d=0.96933.

