## Supplementary Figure 1 for "The matrisome contributes to the increased rigidity of the bovine ovarian cortex and provides a source of new bioengineering tools to investigate ovarian biology"

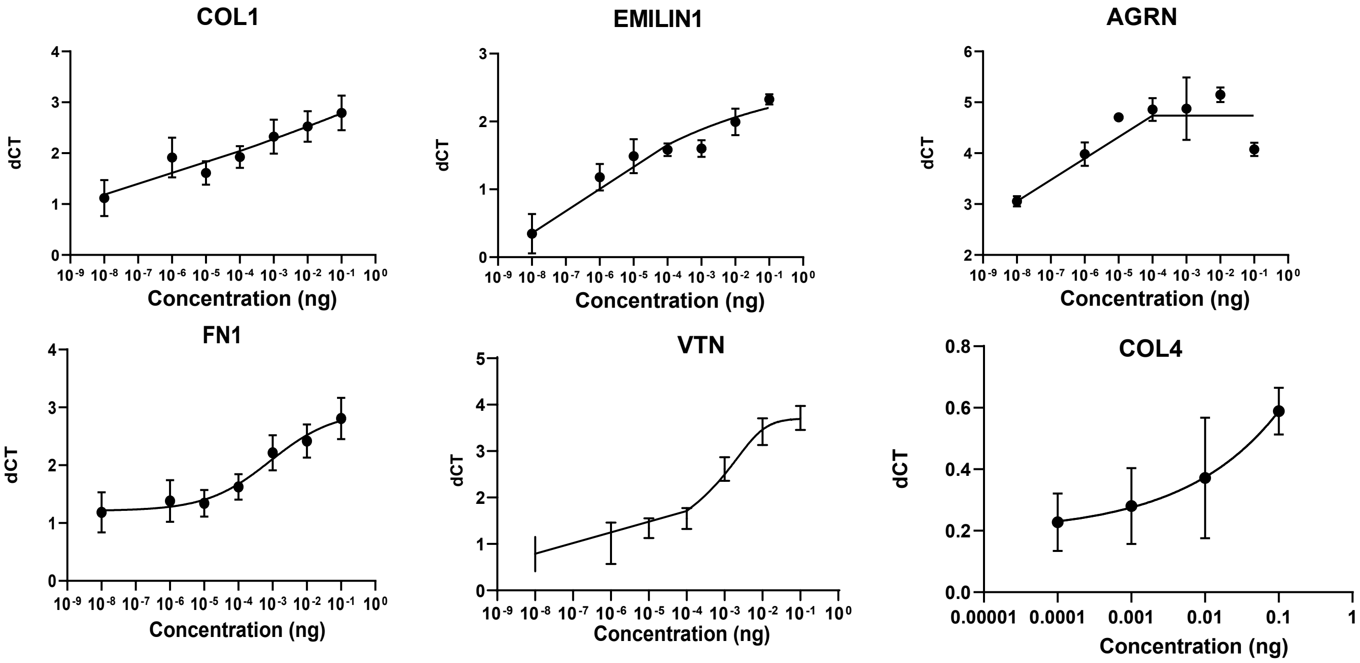


**Standard curves used for interpolation of concentrations of protein.** Bars represented as mean, SEM; N = 3 biological replicates, 2 technical replicates.
