## Supplementary Figure 2 for "The matrisome contributes to the increased rigidity of the bovine ovarian cortex and provides a source of new bioengineering tools to investigate ovarian biology"

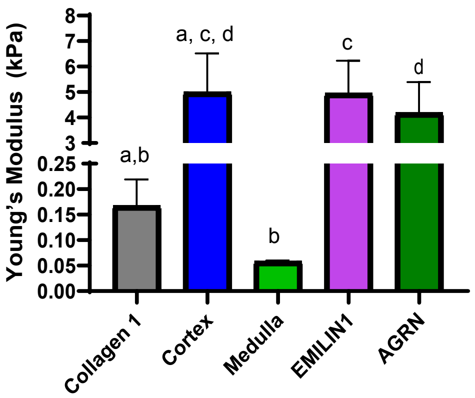


Combined AFM data for FRESH printed scaffolds including collagen 1 only scaffold control. Bars equal to mean and standard error, P-values for relationships: a=0.0196, b=0.0782, c=0.9825, d=0.96933.
