## Supplementary Table 1 for "The matrisome contributes to the increased rigidity of the bovine ovarian cortex and provides a source of new bioengineering tools to investigate ovarian biology"

**Supplementary Table 1:** Antibodies and Proteins used for iPCR and MAPF

| Target | Manufacturer | SKU | Use |
|---|---|---|---|
| AGRN | Abcam | ab85174 | iPCR/MAPF |
| AGRN | LifeSpan Biosciences | LS-G81348-20 | Protein Std |
| COL1 | Abcam | ab90395 | iPCR/MAPF |
| COL1 | EMD Millipore | 234138-1MG | Protein Std |
| EMILIN1 | Fisher Scientific | PA551745 | iPCR/MAPF |
| EMILIN1 | Novus Biologicals | H00011117-P01 | Protein Std |
| COL4A2 | Abclonal | A7657 | iPCR |
| FN1 | Abcam | ab23750 | iPCR |
| VTN | Abcam | ab140016 | iPCR |
